# The phenotype and genotype of fermentative microbes

**DOI:** 10.1101/2023.01.12.523810

**Authors:** Timothy J. Hackmann

## Abstract

Fermentation is a major type of metabolism carried out by many organisms. The study of this metabolism cuts across many fields, including cell biology, animal and human health, and biofuel production. Despite this broad importance, there has been no systematic study of fermentation across many organisms. Here we explore the phenotype and genotype of fermentative prokaryotes in order to gain insight into this metabolism. We assembled a dataset containing phenotypic records of 8,350 organisms (type strains) plus 4,355 genomes and 13.6 million genes. Fermentation was widespread, being found in 30% of all organisms and across the tree of life. Fermentative organisms were more likely than non-fermentative ones to have certain phenotypic traits. Some traits (such as oxygen insensitivity) were expected, but others (such as long cells) were surprising. Fermentative organisms also had a distinct genotype, with 9,450 gene functions and 337 metabolic pathways being more common in them. In a related analysis, we identified end products (metabolites) for 1,455 organisms fermenting 100 substrates. We found 55 products were formed in nearly 300 combinations, showing fermentation is more complex than previously realized. Additionally, we built metabolic models for 406 organisms and predicted which products they form. These models did not predict all products accurately, revealing gaps in our knowledge of metabolic pathways. Our study paints a full picture of fermentation while showing there is still much to learn about this type of metabolism. The microbiology community can explore resources in this work with an interactive tool (https://github.com/thackmann/FermentationExplorer).

## INTRODUCTION

Fermentation is a major type of metabolism. During fermentation, organic molecules (e.g., glucose) are catabolized and donate electrons to other organic molecules. In the process, ATP and organic end products (e.g., lactate) are formed. Because fermentation forms ATP without using O_2_, it is an important alternative to aerobic respiration. The end products are important in human and animal health^1–6^, food^7,8^, and as biofuels or other chemicals^9,10^. Fermentation is thus important in many contexts.

Despite the importance of fermentation, we know little about the range of organisms that carry it out and their traits. There are excellent reviews of fermentation^11–16^, but their focus is on a few model organisms and their biochemical pathways. Information on more organisms is available in journal articles or book chapters^17,18^, but each covers just a few (usually related) organisms. With no central resource for information, it is hard to answer even simple questions about the organisms carrying out fermentation.

Recently, our lab and others have been collecting information on fermentative organisms. Our lab has cataloged which prokaryotes carry out fermentation and which form one end product (acetate)^19,20^. Another group has catalogued end products of fermentation, but the scope is limited to bacteria of the human gut^21^. Other groups have collected information on fermentation when building databases on microbial phenotypes^22,23^. Given the scope of these databases, information specific to fermentation tends to be less complete. Though limited in scope or completeness, these resources provide a good starting point, and if expanded, could give a full picture of fermentative organisms.

Here assemble a large dataset on fermentative prokaryotes and use it to explore the phenotype and genotype of such organisms. This dataset includes phenotypic records on *n* = 8,350 organisms (prokaryotes) as well as *n* = 4,355 genomes, and *n* = 13.6 million genes. With it, we have answered simple but important questions about fermentation. We also built an interactive tool for the microbiology community to explore our dataset and make predictions about organisms of interest.

## RESULTS

### Our dataset contains information on over 8,300 organisms

To examine fermentative organisms, we assembled a dataset from multiple sources (Fig. 1, Supplementary Table 1, Supplementary Table 2). Our starting point was *n* = 1,828 articles from *Bergey’s Manual of Systematic of Archaea and Bacteria*^17^ and the primary literature. From these articles, we obtained names, written descriptions, and phenotypic information for *n* = 8,350 organisms (type strains). The phenotypic information we obtained was for traits related to fermentation (fermentative ability, fermentation end products, fermentation substrates). We used computational approaches to automate some steps (e.g., obtaining names of organisms). Other steps (e.g., obtaining phenotypic information) were done by manually reading articles.

**Fig. 1.**
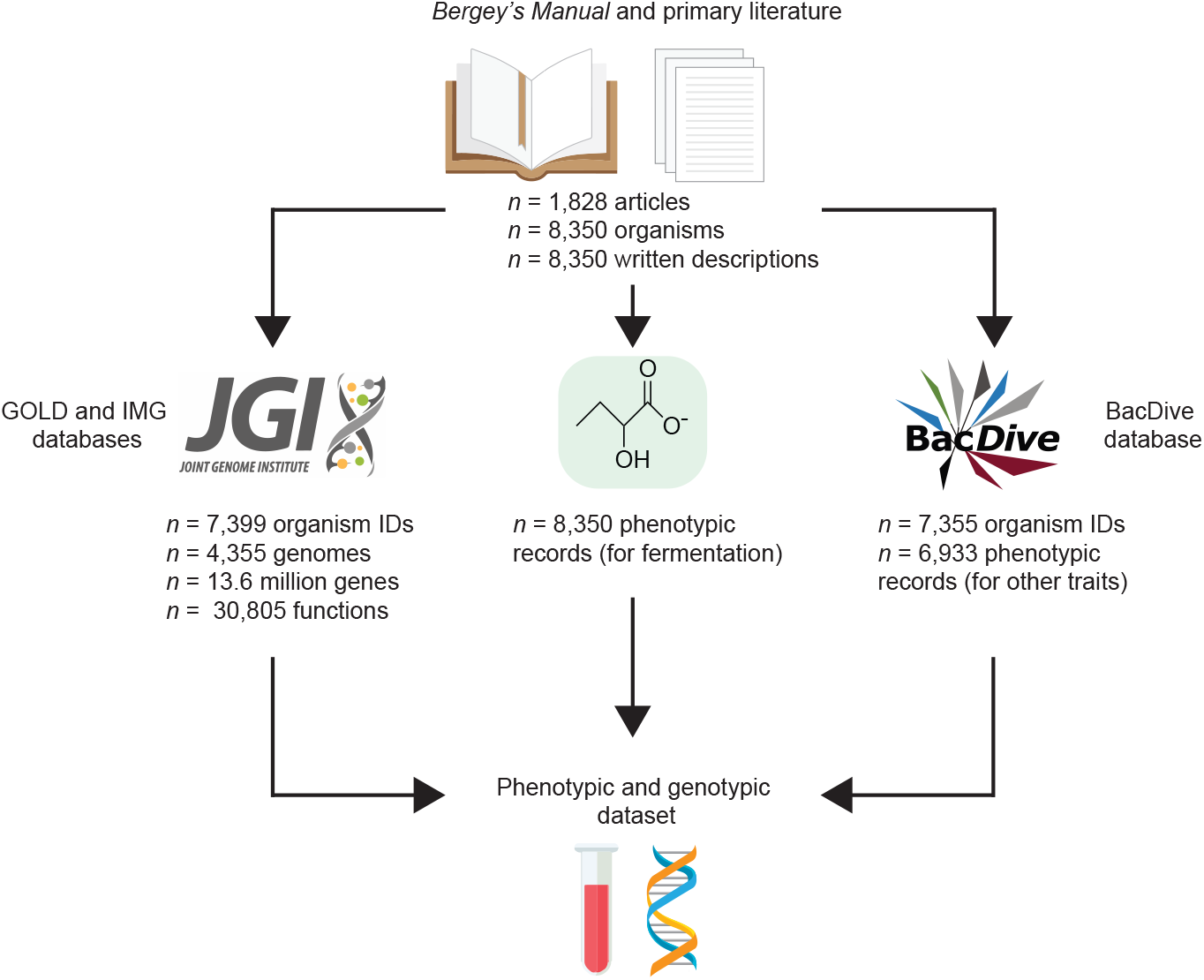
To study fermentative prokaryotes, we assembled a phenotypic and genotypic dataset for *n* = 8,350 organisms (type strains).

We obtained genomic and more phenotypic information from additional sources. We used GOLD^24^ and IMG/M^25^ database to obtain information on genomes and genes from these organisms. We also used HydDB^26^ for information on genes for hydrogenases. Using BacDive database^22^, we obtained phenotypic information for *n* = 14 traits. The traits (e.g., cell length and habitat) were those not directly related to fermentation. Data from BacDive comes from other databases^27^, the primary literature, and unpublished records. We considered other sources of information. For example, we considered one source with data for *n* = 701 organisms of the human gut^21^. However, only *n* = 279 of these were type strains, and *n* = 234 (84%) were already in our dataset. Further, the source used genus-level data when information for specific organisms was not available. Our dataset included only strain-specific data, and we chose not to mix these two different types of information. In sum, our dataset was comprehensive, and it allowed us to probe both the phenotype and genotype of fermentative organisms.

### Fermentative prokaryotes are diverse

With our dataset, we first wanted to answer first how widespread fermentation is across prokaryotes. In total, we found *n* = 2,357 of the 8,350 organisms were capable of fermentation, or 28% of the total (Supplementary Table 1). In earlier work with a smaller dataset^20^, we found a comparable value (33%).

We then built a phylogenetic tree for the *n* = 3,822 organisms with enough information (full ribosomal sequences) available (Fig. 2a). Fermentative organisms were found all over the tree, though they appeared more abundant on some branches that others. The phylum *Firmicutes* was one group in which they were abundant. Non-fermentative prokaryotes were abundant in *Actinobacteria.*

**Fig. 2.**
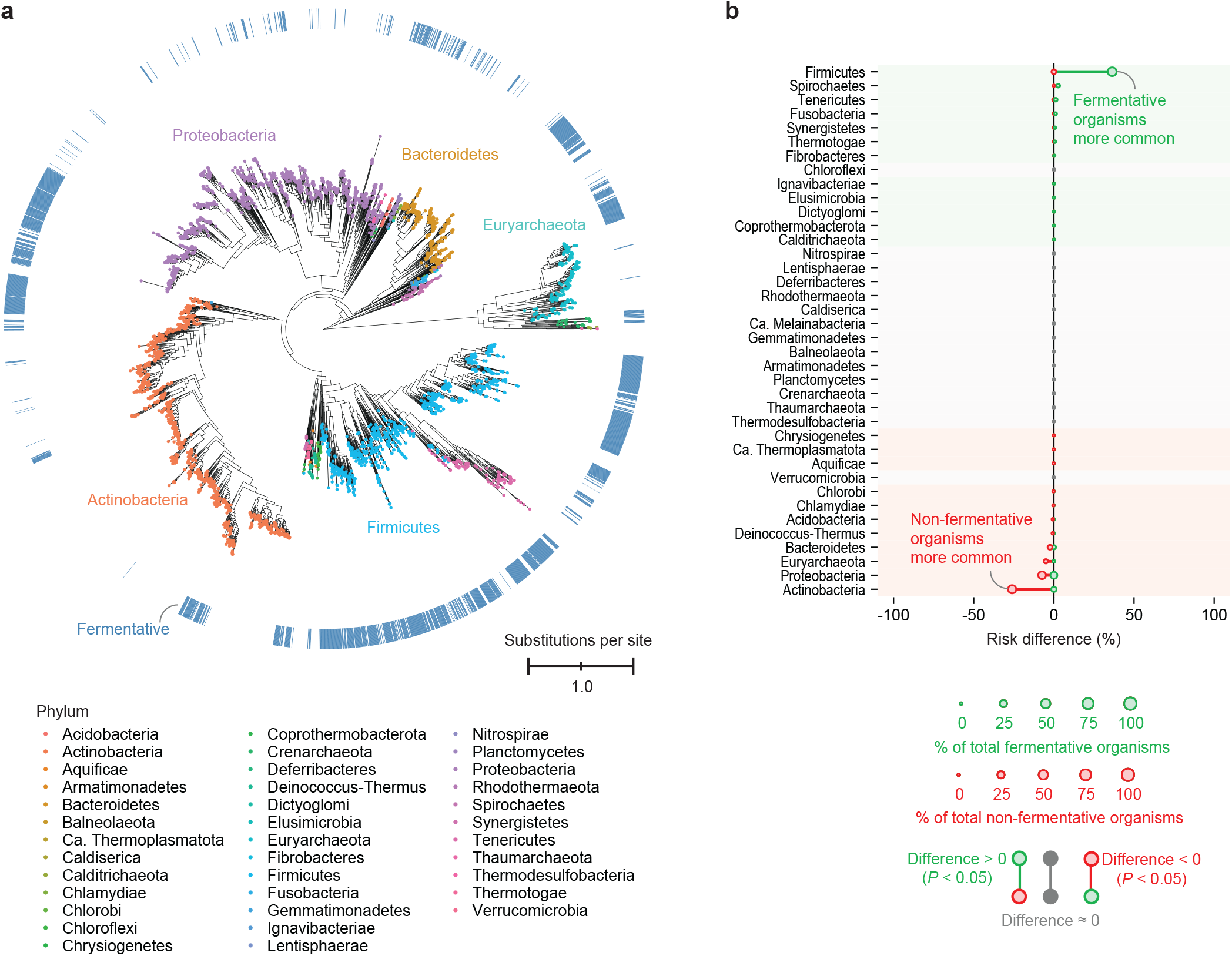
Fermentation is widespread, though it is more common in some groups of organisms than others. **a**, Phylogenetic tree of *n* = 3,822 prokaryotes and their fermentative ability. **b**, Phyla of *n* = 7,395 prokaryotes and their fermentative ability. Names of phyla correspond to the NCBI taxonomy of organisms (see Supplementary Table 1).

We quantified the difference in abundance across phyla using a statistic known as risk difference (see Supplementary Fig. 1a for definition). This analysis was done with *n* = 7,395 organisms. We found the risk difference was highest for *Firmicutes* and lowest for *Actinobacteria* (Fig. 2b), in agreement with the tree (Fig. 2a). It identified *Spirochaetes, Tenericutes,* and *Fusobacteria* as other phyla where fermentative organisms were abundant.

Risk difference is only one example of an effect size statistic. Another commonly used statistic is the log-odds ratio. When we calculate the log-odds ratio, the results are generally similar those with the risk difference (Supplementary Fig. 2). However, many values were undefined (negative or positive infinity), which makes interpretation more difficult.

Together, these results show the ability to ferment is found all over the tree of life, being particularly common for members of *Firmicutes* and certain other phyla.

### Fermentative prokaryotes have distinct phenotypic traits

After identifying which prokaryotes were fermentative, we wanted to answer if their phenotype differed from that of non-fermentative prokaryotes. We examined *n* = 14 types of phenotypic traits, such as habitat, that are not immediately related to fermentation. We calculated the risk difference to determine if traits were more common in fermentative vs. non-fermentative organisms. For continuous traits, we instead calculated a statistic we call the probability difference. The probability difference is simply the difference in the probability density functions between the two groups (see Supplementary Fig. 1b for definition).

We found several traits were more common in fermentative vs. non-fermentative prokaryotes (Fig. 3). Some differences in traits, such as oxygen tolerance, were unsurprising. Fermentative prokaryotes were more likely to be oxygen intolerant (anaerobes or microaerophiles) (Fig. 3a), which follows from fermentation not using O_2_. They were also more likely to be isolated from host-associated habitats (Fig. 3a, Supplementary Fig. 3, Supplementary Fig. 4). Many host-associated habitats, such as the gut, are well known to harbor fermentative organisms^4,28–30^. Some differences in traits were more surprising. For example, fermentative prokaryotes were also more likely to grow with shorter incubation time (Fig. 3b) and have longer cell length (Fig. 3b). Examining the raw data confirms a clear difference in cell length (Supplementary Fig. 5). Though this was surprising, it has been observed that growing cells under microaerophilic or anaerobic conditions increases length^31,32^.

**Fig. 3.**
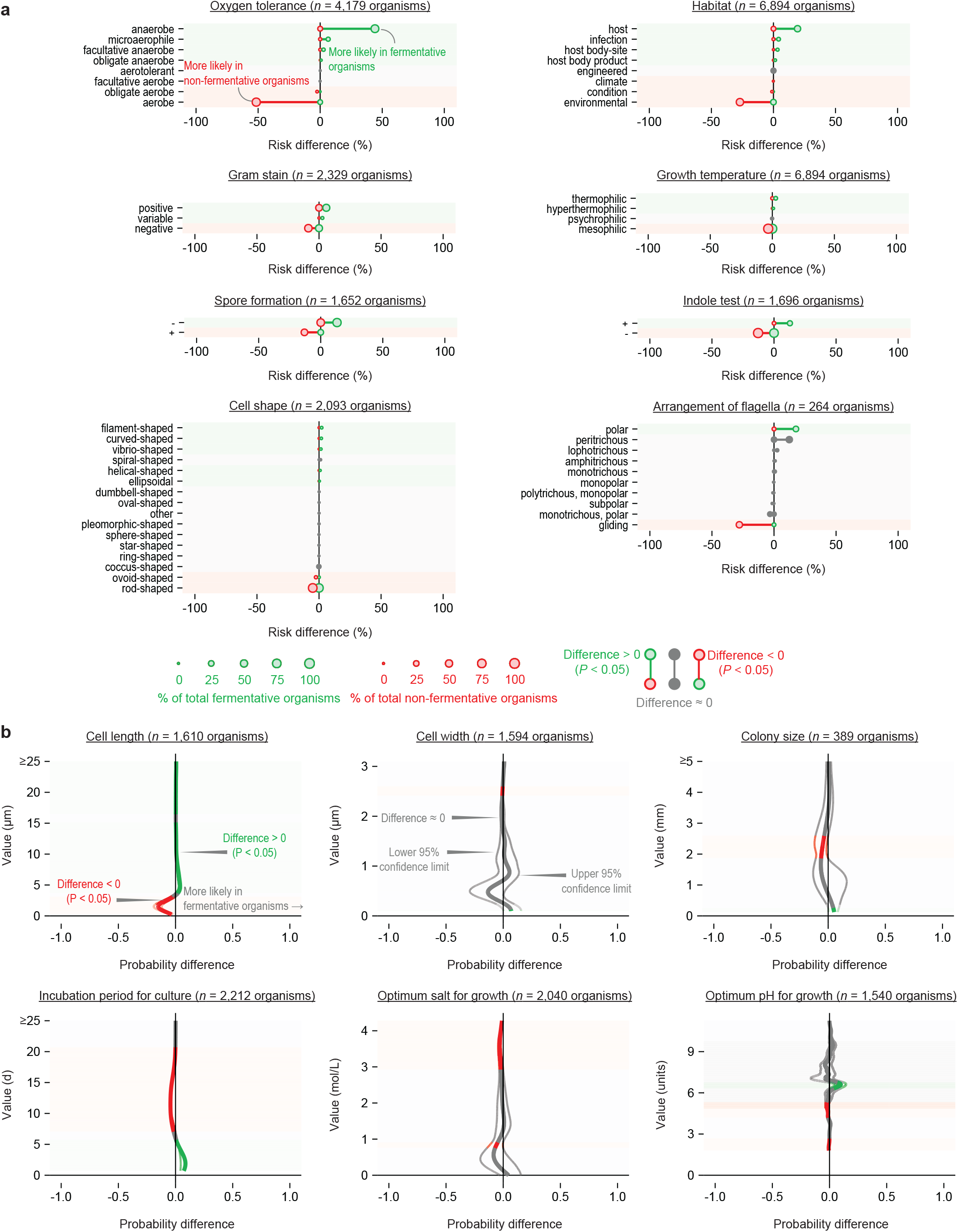
Examining phenotypic records of *n* = 6,933 prokaryotes shows that fermentative organisms have distinct traits. **a**, Discrete traits. The number of organisms per trait is shown in the figure. Habitat refers to the category 1 isolation source. See Supplementary Fig. 3 and 4 for category 2 and category 3 isolation sources. **b**, Continuous traits.

Though the differences were large, they were not as stark as they might be imagined to be. Fermentative prokaryotes were less likely than non-fermentative prokaryotes to be aerobic, but still 18% of them still were (see Fig. 3a). The aerobic, fermentative organisms belonged mostly to *Paenibacillus, Corynebacterium, Staphylococcus, Vibrio, Enterococcus,* and *Streptococcus.* End products and substrates for fermentation have been reported for many of these aerobic organisms (see Supplementary Table 1), confirming they are fermentative.

Our analysis above considered all organisms, and we next examined anaerobic organisms only. We examined a total of *n* = 1,496 anaerobic organisms, *n* = 1,029 of which were fermentative. Even when looking at this subset of data, we still found many traits were more common among fermentative organisms (Supplementary Fig. 6). As before, fermentative organisms were more likely to be isolated from host-associated habitats, grow with shorter incubation time, and have longer cells. Thus, their long cells may owe partly to growth under anaerobic conditions^31,32^, but it is not the full explanation. These results shows that fermentative organisms differ from anaerobic organisms at large.

These results show that fermentative prokaryotes have a distinct phenotype. Some aspects of this phenotype, such as oxygen tolerance, are easy to explain. Others, such as cell length, are surprising but clear. Most fermentative prokaryotes are anaerobic, but many are not, and they still have a phenotype distinct from anaerobic organisms.

### Fermentative prokaryotes have a distinct genotype

Next, we wanted to see if fermentative and non-fermentative prokaryotes differed in genotype. We first looked at their genes and if their predicted functions differed. Our analysis included *n* = 13.6 million genes and *n* = 30,805 predicted functions. These were from *n* = 4,355 organisms, *n* = 1,490 of which are fermentative. The predicted functions correspond to KO^33^, COG^34^, pfam^35^, and TIGRFAM^36^ IDs. We also included functions corresponding to HydDB names^26^.

We found *n* = 9,450 predicted gene functions that were more common in fermentative vs. non-fermentative prokaryotes (Fig. 4a, Supplementary Table 3) (*P* < 0.05). We examined gene functions corresponding to KO IDs first (Fig. 4a). Of the top 15 KO IDs, several are involved production of fermentation end products (e.g., K00656) or their utilization (e.g., K21636). We next looked at COG, pfam, and TIGRFAM IDs. Most predicted functions similar to those of KO IDs, though COG and pfam had more unknown functions (Supplementary Table 3 and below).

**Fig. 4.**
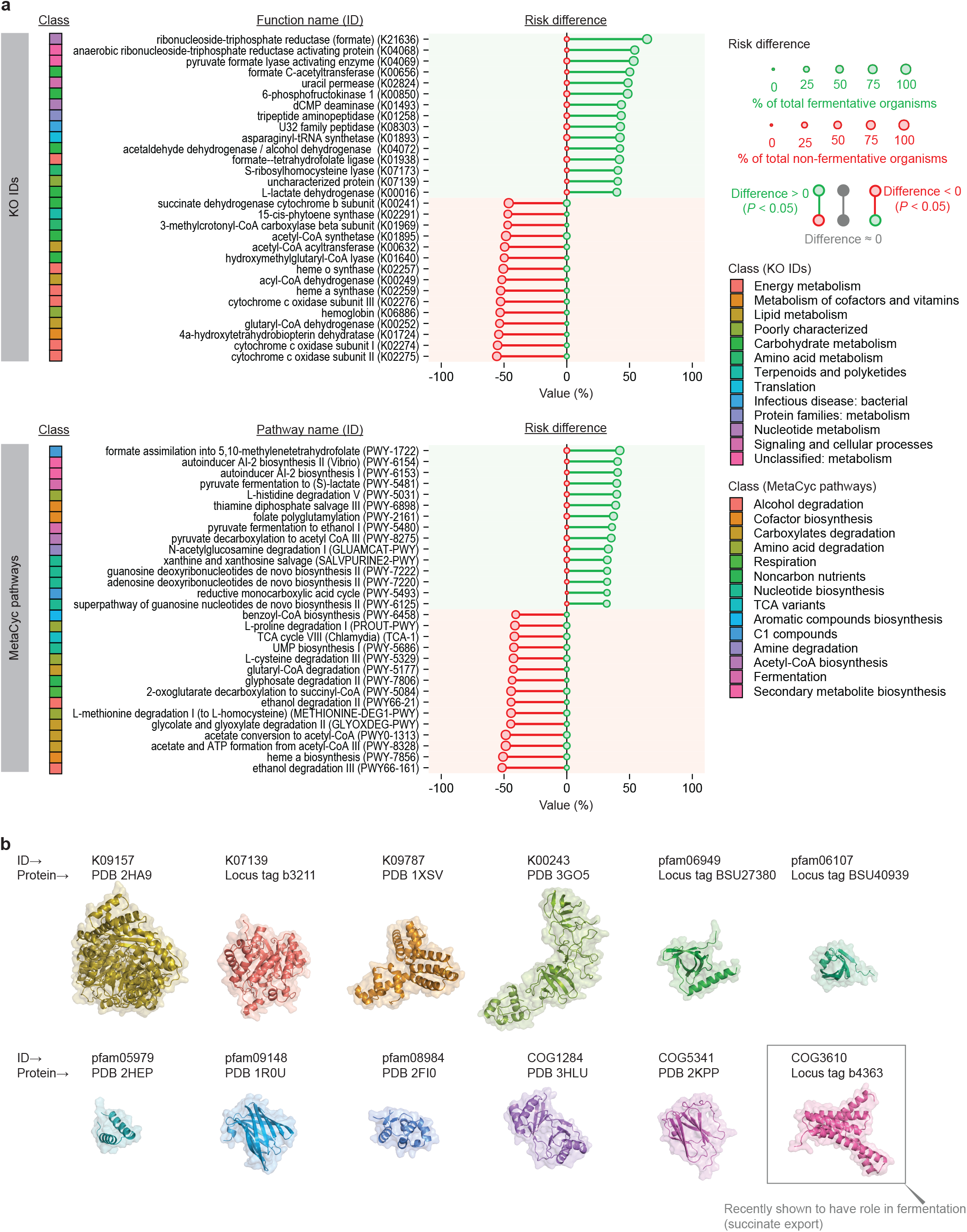
Examining *n* = 13.6 million genes reveals that fermentative organisms have distinct genotype. **a**, Gene functions and predicted MetaCyc pathways. Gene functions for KO IDs alone are shown. See Supplementary Table 3 for full list of gene functions and Supplementary Table 4 for pathways. The class of KO IDs refers to level B of the BRITE hierarchy. The class for MetaCyc pathways refers to the second-highest parent class. Some class names have been abbreviated for display. **b**, Structures of uncharacterized proteins common in fermentative organisms.

Most gene functions were clear and well defined, but many were not. In total, *n* = 1,229 gene functions more common in fermentative organisms were “uncharacterized” or of “unknown function” (Supplementary Table 3). We found or predicted protein structures for several of these (Fig. 4b). Despite having no established function, they would appear to be important role in fermentation. In one case, this has been verified; COG3610 is still reported as uncharacterized in its database, but one member was recently characterized as a transporter of a fermentation product (succinate)^37^.

We also found *n* = 13,945 gene functions that were more common in non-fermentative prokaryotes (Fig. 4a, Supplementary Table 3) (*P* < 0.05). Many corresponded to enzymes used in oxidative phosphorylation (e.g., K02275).

Besides looking at the level of individual genes and functions, we examined if certain metabolic pathways were more common. To do this, we examined the abundance of all *n* = 3,445 metabolic pathways in MetaCyc^38^ (Fig. 4a, Supplementary Table 4). The prokaryotes in our dataset encoded *n* = 1,579 of these pathways, and *n* = 337 were more abundant in fermentative ones (*P* < 0.05). The top 15 pathways in fermentative prokaryotes included those for production of fermentation products (e.g., PWY-5481) or their utilization (e.g., PWY-1722). More surprisingly, there were pathways involved in quorum sensing (e.g., PWY-6154) and nucleotide metabolism. Thus, pathways were as expected, though a few surprises were also present.

Our analysis above considered all organisms, and we next examined anaerobic organisms only. We examined a total of *n* = 1,039 anaerobic organisms, *n* = 745 of which were fermentative. Again, we found gene functions and pathways more common in fermentative organisms (Supplementary Table 5 and 6). Further, they were similar to functions and pathways identified above. Of the top 15 KO IDs in Fig. 4a, 13 were still more common in fermentative vs. non-fermentative organisms (see Supplementary Table 5). The situation was similar for MetaCyc pathways, with 11 of the 15 top pathways in Fig. 4a still being more common (see Supplementary Table 6). These results shows the genotype of fermentative organisms differs from anaerobic organisms at large.

These results together show that fermentative prokaryotes have a distinct genotype. Intriguingly, several genes appear to be important in fermentation but have no known function. These are targets for further study in understanding fermentation and protein function.

### Fermentation produces many end products

Another important trait of fermentative prokaryotes is which end products of fermentation they form. We collected information on end products for *n* = 1,455 organisms. Based on the text of the written description, we divided these into major and minor end products. We also recorded the *n* = 100 substrates reported to form these end products (information available for *n* = 1,260 organisms).

In total, we found prokaryotes formed *n* =55 fermentation products (Fig. 5a,b). Acetate and lactate were the most common products, with at least one being formed by 97% of organisms. Most (83%) organisms formed multiple organic products (Fig. 5c). Of the organisms that formed only one organic product, nearly all formed lactate (56%) or acetate (34%). There were many (*n* = 289) unique combinations of products altogether.

**Fig. 5.**
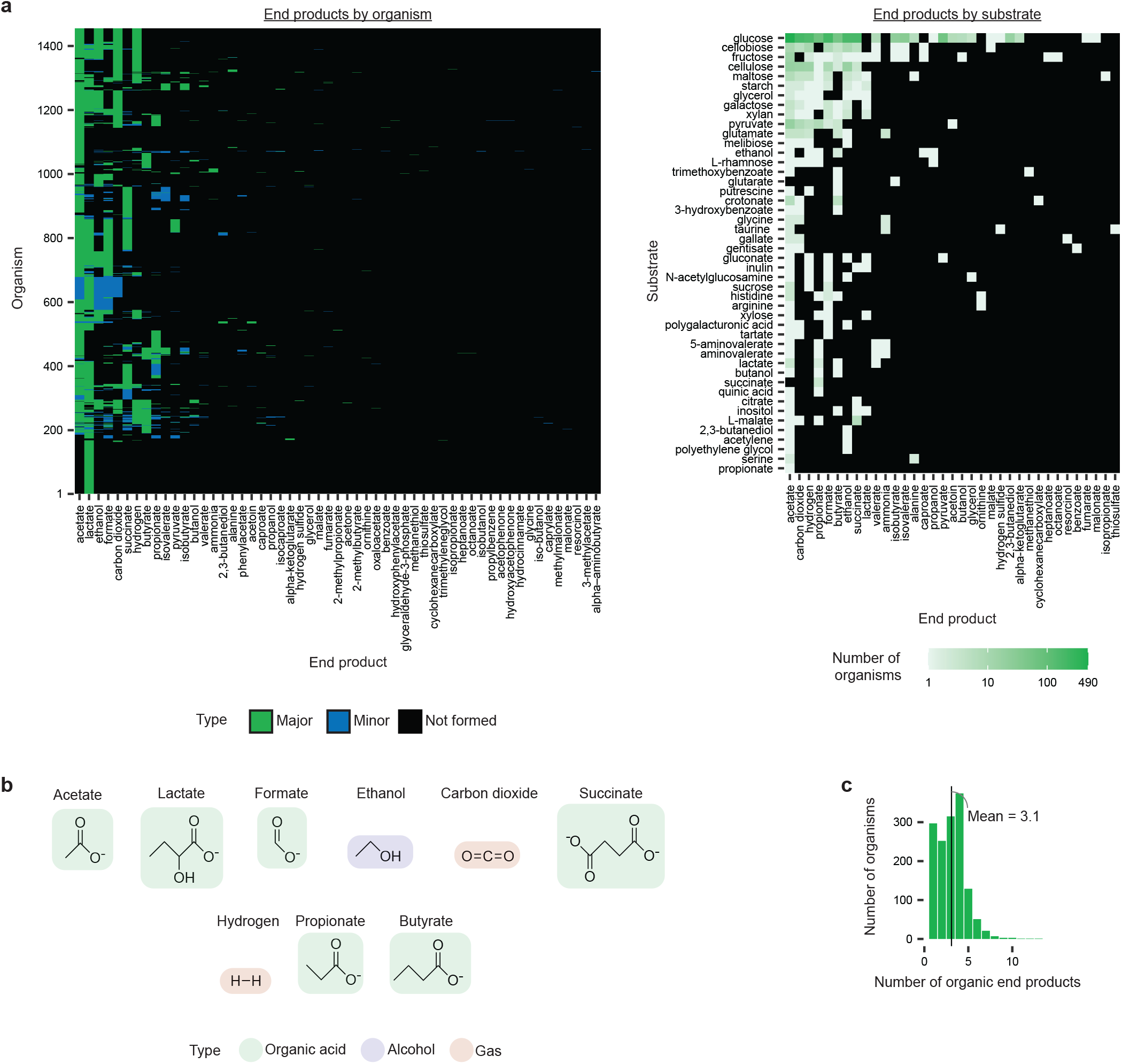
Examining end products of *n* = 1,455 prokaryotes shows that a wide number and range are formed. **a**, Range of products across organisms and substrates. Lactate represents two potentially different products [(S) lactate and (R) lactate], but articles did not always distinguish which was formed. **b**, Most common fermentation products. **c**, Number of products formed.

We also examined end products formed by substrate (Fig. 5a,b). We focused on the *n* = 46 substrates that are single, chemically-defined molecules (e.g., glucose). This represented information for *n* = 805 organisms. Glucose was the most common substrate reported, and acetate was the most common product (Fig. 5a). Other common products were as before, though the order of their abundance differed somewhat (Fig. 5a,b).

The number of end products in our dataset is higher than in other sources. It is >3-fold higher than reported for prokaryotes of the human gut^21^ (Supplementary Fig. 7). The large size of our dataset makes it a useful resource.

Our dataset gives simple but important insights about end products of fermentation. It shows which products are most common, and it shows nearly all fermentations form multiple products and in many combinations. It shows the relationship between substrate, organism, and end product with a dataset of unprecedented size.

### End products can be predicted from genes using metabolic models

Many studies use an organism’s genes to predict their metabolic pathways and fermentation products they form^21,39–41^. We wanted to test how reliable is this practice with our dataset. We built metabolic models of our organisms, then used flux balance analysis^42^ to predict end products. We then compared predicted end products to those observed in our dataset. Because glucose metabolism is well studied, we examined organisms known to ferment glucose (*n* = 406) and focused on the most *n* = 9 most common fermentation end products (see Fig. 5b).

The metabolic models we built were networks of up to *n* = 98 metabolites connected by *n* = 110 biochemical reactions and *n* = 209 enzymes (Fig. 6a). Of the total metabolites, *n* = 64 were carbon-carrying metabolites (e.g., glucose), and an additional *n* = 34 were cofactors (e.g., NADH). For each organism, we built a model containing only those enzymes and reactions corresponding to the KO IDs of its genes (see Fig. 6b). For hydrogen formation, we used reactions corresponding to the HydDB name instead. With each model, we calculated the flux (flow) of metabolites through the network. If there was a positive flux from the substrate (glucose) to the endproduct (e.g., lactate), that endproduct was predicted to be formed (see Fig. 6b).

**Fig. 6.**
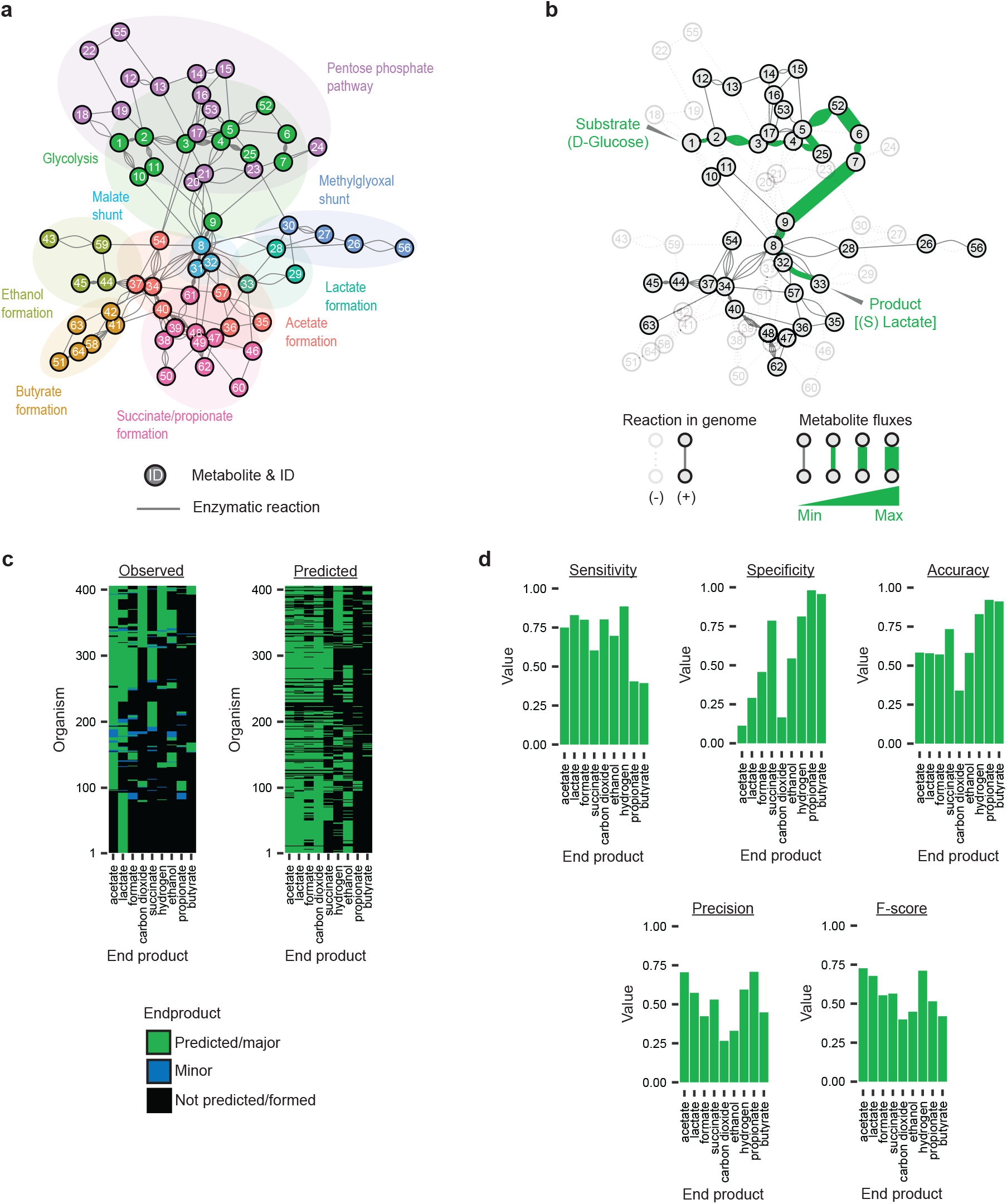
Metabolic models of *n* = 406 organisms predict end products of fermentation. **a**, The reference model containing and *n* = 209 enzymes, *n* = 110 biochemical reactions, and *n* = 64 metabolites (cofactors excluded). **b**, An example model for one organism *(Lactococcus plantarum),* showing the predicted flux from D-glucose to (S)-lactate. **c**, End products predicted by metabolic models compared to those observed in our dataset. **d**, Performance of predictions. In (c), the metabolites are 1. D-glucose, 2. D-glucose 6-phosphate, 3. D-fructose 6-phosphate, 4. D-fructose 1,6-bisphosphate, 5. D-glyceraldehyde 3-phosphate, 6. 3-phospho-D-glycerate, 7. 2-phospho-D-glycerate, 8. pyruvate, 9. phosphoenolpyruvate, 10. protein histidine, 11. protein n(pi)-phospho-l-histidine, 12. D-glucono-1,5-lactone 6-phosphate, 13. 6-phospho-D-gluconate, 14. D-ribulose 5-phosphate, 15. D-ribose 5-phosphate, 16. sedoheptulose 7-phosphate, 17. D-xylulose 5-phosphate, 18. D-glucono-1,5-lactone, 19. D-gluconic acid, 20. 2-dehydro-3-deoxy-D-gluconate, 21. 2-dehydro-3-deoxy-6-phospho-D-gluconate, 22. 2-keto-D-gluconic acid, 23. D-glyceraldehyde, 24. D-glycerate, 25. glycerone phosphate, 26. (R)-S-lactoylglutathione, 27. (R)-lactaldehyde, 28. (R)-lactate, 29. (S)-lactaldehyde, 30. methylglyoxal, 31. (S)-malate, 32. oxaloacetate, 33. (S)-lactate, 34. acetyl-CoA, 35. enzyme N6-(lipoyl)lysine, 36. enzyme N6-(dihydrolipoyl)lysine, 37. acetate, 38. succinyl-CoA, 39. succinate, 40. CoA, 41. butanoyl-CoA, 42. butanoic acid, 43. thiamin diphosphate, 44. acetaldehyde, 45. ethanol, 46. lactoyl-CoA, 47. propanoate, 48. propanoyl-CoA, 49. (S)-methylmalonyl-CoA, 50. (R)-methylmalonyl-CoA, 51. (S)-3-hydroxybutanoyl-CoA, 52. 3-phospho-D-glyceroyl phosphate, 53. D-erythrose 4-phosphate, 54. acetyl phosphate, 55. 6-phospho-2-dehydro-D-gluconate, 56. glutathione, 57. [dihydrolipoyllysine-residue acetyltransferase] s-acetyldihydrolipoyllysine, 58. butanoylphosphate, 59. 2-(alpha-hydroxyethyl)thiamine diphosphate, 60. propenoyl-CoA, 61. fumarate, 62. propanoyl phosphate, 63. acetoacetyl-CoA, and 64. crotonoyl-CoA.

We compared products predicted by models with the observed ones (Fig. 6c). Some products (acetate, lactate, formate, CO_2_) were predicted in nearly all organisms, whether they were observed to be formed or not. This meant the predictions had high sensitivity, but low specificity (Fig. 6d). Other products (propionate, butyrate) were predicted in far fewer organisms than were observed to form them (Fig. 6c). The predictions had high specificity, but low sensitivity (Fig. 6d). Three products (succinate, ethanol, hydrogen) were predicted with high sensitivity and specificity (Fig. 6d).

We explored why certain products were predicted with low specificity (i.e., had many false negative predictions). For butyrate, we were able to identify a missing reaction in 45% of cases (Supplementary Fig. 8). The most common reaction missing was butyryl-CoA:acetate-CoA transferase (EC 2.8.3.8), and adding it back to an organism’s model restored its ability to produce butyrate. This reaction may be missing because of poor gene annotation—databases such as KEGG are missing 86% of enzymes known to carry out this reaction^43^. In 20% of cases, more than one reaction could restore the ability to produce butyrate, making it unclear which reaction was really missing. In the remaining cases, the missing reaction(s) could not be identified. For propionate, another product predicted with low specificity, the situation was similar (Supplementary Fig. 8).

Our study shows that some, but not all, end products could be reliably predicted from metabolic models. Some products were predicted with low specificity, indicating more organisms have the ability to produce than observed (or reported). Others were predicted with low sensitivity instead. Further, our study shows a number of reactions apparently missing from organisms. Poor gene annotation is likely to blame in some cases, though the reason is not clear in others. Our study shows that are still gaps in our knowledge of metabolic pathways and their prediction, even for glucose.

### Our work is available in an interactive tool

To maximize the usefulness of our work, we built an interactive tool we call Fermentation Explorer (Fig. 7). This tool allows users to explore our dataset and make predictions about organisms in their samples.

**Fig. 7.**
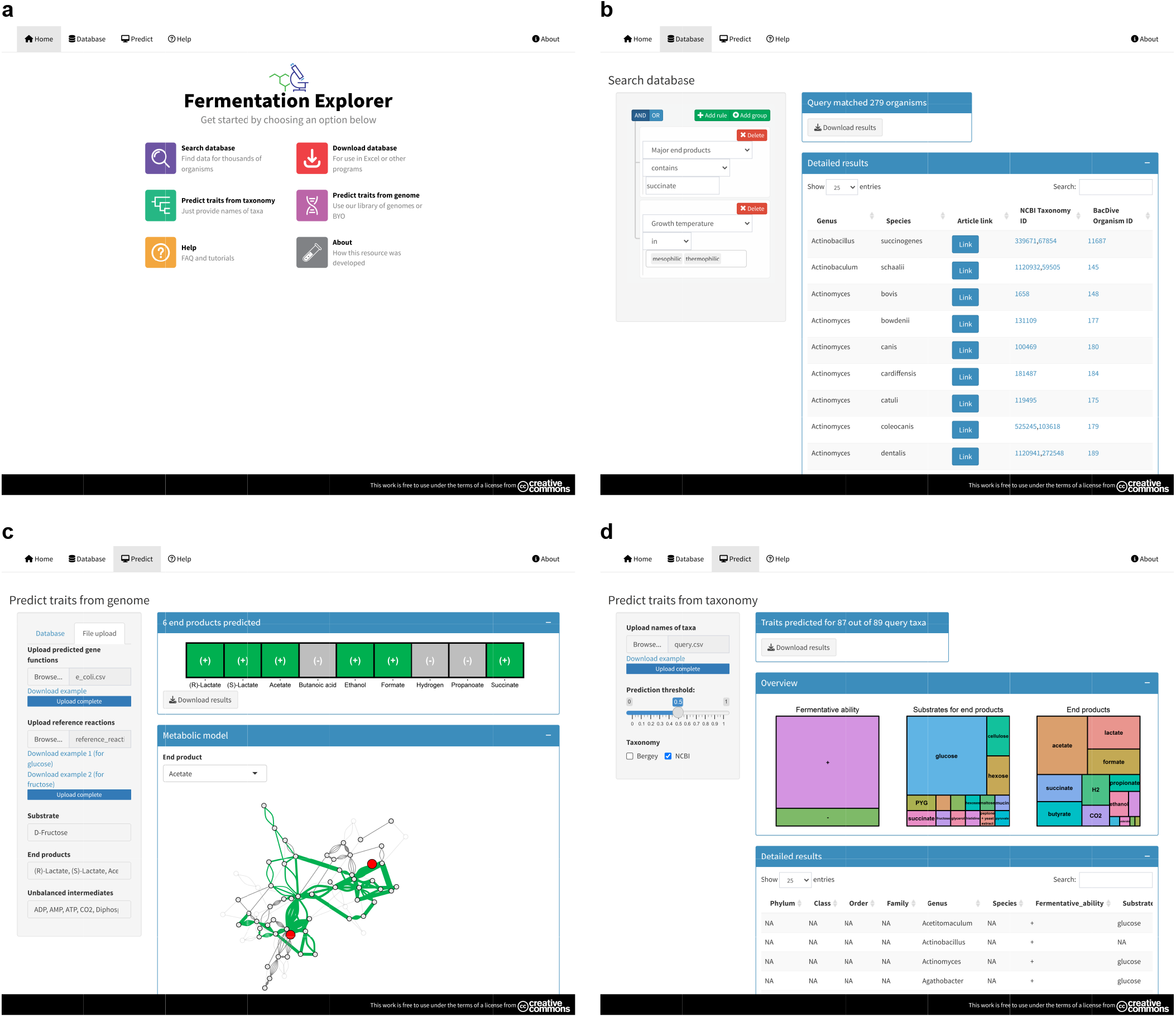
Screenshot of Fermentation Explorer, an interactive tool for exploring our dataset and making predictions about organisms. **a**, Home page. **b**, Database search. **c**, Predicting end products with metabolic models. **d**, Predicting fermentative traits from taxonomy.

With this tool, users can build metabolic models and predict fermentation end products for their organisms (Fig. 7c). Users first upload KO IDs for genes for an organism. After this, they specify substrates and end products to check. The tool builds a metabolic model specific to the organism and predicts products formed. Users can also visualize the model with predicted fluxes. To provide KO IDs, users can use the KEGG Automatic Annotation Server^44^—the output can be uploaded into our tool. Users can also use our library of KO IDs for *n* = 987 organisms (preloaded into the tool).

Users can also predict traits for organisms from their taxonomy (Fig. 7d). After the user uploads the taxonomy (names) of their organisms, the tool finds matching organisms from our dataset. If the matching organisms share a trait, the trait is predicted for the user’s organisms. By default, all matching organisms must share the trait, but this threshold is adjustable by the user. The method of prediction is similar to FAPROTAX^23^, but with this adjustable threshold added. The tool accepts partial taxonomy (e.g., resolved at genus only), making it useful for organisms identified with ribosomal DNA sequencing. The user can also choose different systems for taxonomy (that from *Bergey’s Manual* or NCBI).

The tool is available for use online and for download (https://github.com/thackmann/FermentationExplorer). It is built as an R Shiny app, and the online version is hosted by shinyapps.io. The app can be downloaded for use within R or as a Docker container image. Being easy to access and use, the tool enables the microbial community to apply the resources in our work in their own research.

## DISCUSSION

Fermentation is a major type of metabolism, and it is important in microbial physiology, host health, food production, and industry. Despite this importance, there has been no systematic study of fermentative organisms and their properties. Reviews of fermentation have been based on information from model organisms^11–16^, which may not capture the full diversity of this metabolism. Some work, including our own, had started to accumulate information on more organisms^19,20,22,23^, but a full picture has still been lacking.

By using a dataset of *n* = 8,350 organisms (28% of which are fermentative), the current study paints a fuller picture. It shows that fermentative organisms are both abundant and widespread. It reveals key insights into their genotype and phenotype and thus the concept of fermentation as a whole.

Some of the insights we reach make sense in context of earlier work, while others are surprising. Some are both expected and surprising at the same time. Given that fermentation does not use oxygen, it is unsurprising that many organisms were reported as anaerobes. Indeed, fermentation was first described as “la vie sans air” (life without air)^45^, and it is still regarded as a principally anaerobic metabolism^11–16^. It is thus all the more surprising that 18% of fermentative organisms were reported as aerobes. These organisms may be more oxygen tolerant than previously realized.

One important insight is fermentation is complex and forms many end products. Historically, fermentations have been named by the major end product they form—e.g., alcohol fermentation forms ethanol^11–13,15,16^. This is a long-standing practice^46,47^ and remains useful for teaching, but our dataset shows it does not match reality. Nearly all fermentations formed multiple products and in nearly 300 combinations. Our work reveals a complexity to fermentation not fully apparent before.

With the rise of genome sequencing, it is common to use an organism’s genome to predict its metabolic pathways and fermentation products^21,39–41^. This practice is especially important when an organism is uncultured and known only by a genome sequence^40,41^. Our study shows this practice is useful but has limits. We used genomes of *n* = 406 organisms to build metabolic models of fermentation. Some end products were predicted with low specificity or low sensitivity. Low specificity means that more organisms are predicted to form a product than observed. It could be due to many factors, such as a product being formed only under certain growth conditions (often the case for lactate^48,49^). Alternatively, the product could be used for anabolism (often the case for formate^50,51^) and not leave the cell. Low sensitivity means that fewer organisms are predicted to form a product than observed, which represents a different issue.

One approach for raising sensitivity of predictions is gap filling, or adding enzymatic reactions apparently missing from models^52^. This approach is common and used by a study predicting end products for bacteria from the human gut^21^. Our study identified reactions that are often missing, and adding these would have raised sensitivity. However, it was not always clear which reaction was missing, much less the reason. If anything else, missing reactions show areas of metabolism needing further study; such study has led to new pathways for fermentation being discovered^19^.

To maximize the value of our work to the microbial community, we built an interactive tool called Fermentation Explorer. This tool has many applications. One is for identifying organisms for producing end products for biotechnological purposes (e.g., biofuel production). With data on 55 fermentation end products formed in nearly 300 combinations, our tool can pinpoint the best organism for an application. Such an organism could be used outright, or it could be used as a source of genes to genetically engineer other organisms^53–56^. Another application is predicting fermentative traits of organisms present in a user’s sample. Users can predict traits in two different ways—either from an organism’s genome or its taxonomy. Because fermentation is widespread, this ability will be useful to microbiologists working in many systems.

Fermentation has been studied for 185 years (since the time of Theodor Schwann^57^ and Louis Pasteur^46,47^), and our study fills key gaps in our knowledge of this metabolism. It also shows there is much to learn. It shows common genes in fermentative organisms and that some genes have no defined function. The latter are targets for further study, and their abundance in fermentative organisms may help narrow down possible roles. In one case, a gene with no defined function was confirmed to have a role in fermentation (transport of succinate)^37^. Our study shows a wealth of information exists on fermentative prokaryotes, but it speaks little on eukaryotes. Both unicellular and multicellular eukaryotes carry out fermentation^14^, and they merit further study. The next 185 years of study will be illuminating.

## METHODS

### *Collection of information from* Bergey’s Manual *and the primary literature*

We collected information from *Bergey’s Manual of Systematic of Archaea and Bacteria* using an approach similar to ref. ^19,20,58^. We downloaded all *n* = 1,751 articles for genera. We then extracted name, strain ID(s), and written description of each of the *n* = 8,331 organisms using R scripts.

To obtain phenotypic traits related to fermentation, we read all *n* = 5,465 written descriptions containing the keyword “ferment”. The traits we recorded were fermentative ability, fermentation end products, and the substrate used to form those end products. We also recorded the original text reporting the end products, then we used this text to divide fermentation end products into major and minor types. Minor products were those produced in only small quantities or only under certain conditions.

To check information in *Bergey’s Manual,* we found *n* = 77 articles in the primary literature on prokaryotes of the rumen, an environment that our lab studies. These articles^59–135^ cover all *n* = 88 type strains from this environment. We found these articles using an approach similar to ref. ^20,39^. We recorded the types of information above, plus yield of fermentation end products (mol/mol substrate used) in the *n* = 50 cases it was available. For polymers of hexose (e.g., cellulose), yield was expressed as mol/mol hexose equivalents. The moles of substrate used was calculated from moles of carbon in products. When yields were available, minor products were defined as those with yield <0.05 mol/mol.

We compared information in *Bergey’s Manual* and the primary literature, and we found good agreement between the sources (Supplementary Fig. 9, Supplementary Table 2). In total, 78% of organisms in the primary literature were present in *Bergey’s Manual,* and those absent tended to be recently described. Fermentative ability and fermentation end products also closely agreed. For organisms described both in *Bergey’s Manual* and the primary literature, we used information from the latter in subsequent analysis.

### Collection of information from other sources

We obtained information on genomes and genes from GOLD^24^ and IMG/M^25^ database. Following ref. ^19,20,58^, we searched GOLD database for a GOLD organism ID, GOLD project ID, and IMG genome ID for each organism. Organisms were matched to GOLD organism ID by their strain ID plus genus or species name. If the strain ID was assigned by a large culture collection (DSMZ, ATCC, JCM, or CIP), only the strain ID had to match. We then searched IMG/M database for all protein-coding genes belonging to each genome (IMG genome ID). If an organism had multiple genomes, we chose the one with the most genes. For each gene, we downloaded the locus tag as well as gene, KO, COG, pfam, and TIGRFAM IDs.

We obtained phenotypic traits (those not related to fermentation) from BacDive^22^. We searched for BacDive IDs for organisms, matching them by strain ID plus the genus or species name (see above). We downloaded information for traits using the “Advanced search” and “Isolation sources” features. We formatted data as needed, such as by averaging data given as ranges (e.g., pH 7-9). Unformatted data are presented in Supplementary Table 1.

We classified putative genes for hydrogenases using HydDB^26^. We obtained sequences of genes using IMG/M and database IDs (pfam02906, pfam00374, TIGR03295) covering hydrogenases of interest. We then classified sequences with HydDB^26^. HydDB classified the sequences as [NiFe] Group 3b, [NiFe] Group 3d, [NiFe] Group 4a, [FeFe] Group A, or other. We further classified [FeFe] Group A hydrogenases as Group A1, A2, A3, or A4 according to which accessory proteins were adjacent^26^. Members of Group A2 had HytE1 (K05796) adjacent; Group A3 had HydB (pfam01512) adjacent; Group A4 had GltA (COG0493) adjacent; and Group A1 had no accessory protein adjacent.

We obtained definitions (short descriptions) of KO IDs from KEGG^33^, COG IDs from COG^34^, and TIGRFAM IDs^36^ and pfam IDs from InterPro^35^. We obtained NCBI taxonomy ID and names from GOLD database.

We obtained information about organisms in ref. ^21^ by downloading it at https://www.vmh.life/#microbes/fermcarb. We determined which organisms were type strains by using BacDive. We found BacDive IDs for the organisms in that source following methods above.

### MetaCyc pathways

We predicted which of the *n* = 3,445 pathways on MetaCyc^38^ were encoded by genomes. We navigated to the MetaCyc webpage for each pathway, then we downloaded enzyme commission (EC) numbers for enzymes of each pathway using “Download Genes”. We then found a KO ID corresponding to each EC number on KEGG^33^.

We defined a genome as encoding a MetaCyc pathway if it had KO IDs for each enzyme. If an EC number had multiple KO IDs, only one had to match. If an enzyme had no EC number or no KO ID, it was ignored.

### Construction of metabolic models

We constructed metabolic models for organisms using R and the fbar package. We first constructed a reference model that contained all *n* = 110 biochemical reactions we found relevant to fermentation (see Supplementary Table 7). The reactions were identified mostly from ref. ^39^ and are catalyzed by *n* = 209 enzymes. We obtained information on each reaction (including equation, EC number, and KO ID) from KEGG^33^ and other sources. For reactions catalyzed by hydrogenases, we list the HydDB name in place of the KO ID. Many reactions could be catalyzed by multiple enzymes, and many enzymes had multiple KO IDs. Our model preserves the relationship between reaction, enzyme, and KO ID (see Supplementary Table 7). After defining the main reactions, we added exchange reactions representing entry of cofactors, substrates, and products.

We predicted which reactions are catalyzed by each organism. We predict that a reaction is catalyzed if an organism has a gene with the appropriate KO ID (or HydDB name). If a reaction is catalyzed by an enzyme with multiple KO IDs, genes for all must be present for it to be predicted.

Lastly, we set constraints for and solved the model. Fluxes for most reactions were constrained between −1000 and 1000 (arbitrary units). Fluxes for reactions that usually proceed in the forward direction (e.g, EC 2.7.1.1) were constrained between 0 and 1000. Similarly, fluxes for reactions proceeding in reverse were constrained between −1000 and 0. Fluxes of exchange reactions were constrained to be between −10^6^ and 10^6^ (for cofactors), −1000 and 0 (for substrates), or 0 and 10^6^ (for products). For reactions not predicted to be catalyzed, fluxes were constrained to 0. The model was solved once for each product; this was done by setting the objective function of its exchange reaction to be 1. Products with flux >1 were considered to be produced.

Cofactors included molecules such as NADH and ATP. They also included any metabolite, such as H_2_O, that we wished to be unbalanced. They did not include molecules, such as CoA, that are balanced by reactions in close proximity. With cofactors (unbalanced metabolites) in the model, the structure could be much simpler and did not have to include reactions for anabolism.

For propionate and butyrate, we explored how adding reactions would affect predictions. We built models of organisms observed but not predicted to form propionate or butyrate. We then added one-by-one all *n* = 110 biochemical reactions in the reference model. We recorded the flux and determined if adding a reaction would change it from a negative prediction (flux <1) to a positive one (flux >1).

Models were plotted using R and the igraph package^136^.

### Construction of phylogenetic trees

Phylogenetic trees were constructed as in ref. ^19,20^. The construction used sequences of 14 ribosomal proteins^137^. We downloaded sequences from IMG/M^25^, then aligned and concatenated them in R. We used aligned and concatenated sequences to create a phylogenetic tree with RAxML^138^. Branch lengths of the consensus tree were calculated using R and the phytools package^139^. The consensus tree was visualized using R and the ggtree package^140^.

### Protein structures

Structures were downloaded from RCSB PDB^141^ or predicted with ColabFold^142^. Structures on PDB were found using by using COG^34^ and InterPro^35^ as references. If none existed for a given database ID, we predicted structures with ColabFold for *Escherichia coli* or *Bacillus subtilis.* The protein sequence used for prediction was downloaded from IMG/M. Structures were visualized using PyMOL (v. 2.0, Schrödinger, LLC) following ref. ^43^.

### Other bioinformatics

We constructed heatmaps, lollipop charts, and bar charts using R and the ggplot2 package^143^. Our interactive tool was constructed using R and the Shiny package.

### Statistics

We calculated risk difference for discrete variables as

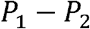

where *P_1_* is the percentage of fermentative organisms positive for a trait and *P_2_* is the corresponding value for non-fermentative organisms (see Supplementary Fig. 1a). We tested if the value was different from 0 using a two-tailed *z*-test^144^. Values of *P_1_* and *P_2_* were arcsine transformed^144^ before the *z*-test, and they were untransformed for presentation in figures. *P*-values from the *Z*-test were corrected for multiple comparisons using the Benjamini–Hochberg procedure^145^.

For continuous variables, we calculated what we call the probability difference (see Supplementary Fig. 1b). It is

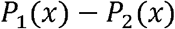

where *P*_1_(*x*) is the probability density function for fermentative organisms. *P*_2_(*x*) is the corresponding function for non-fermentative organisms. We fitted probability density functions to data using the density function of R (setting the bandwidth to triple the default value). We calculated 95% confidence limits using bootstrapping with 10,000 replicates.

## Data availability

All data are available within the paper, supplementary information, and interactive tool.

## Code availability

Code for the interactive tool is available at https://github.com/thackmann/FermentationExplorer.

## Acknowledgements

This work was supported by an Agriculture and Food Research Initiative Competitive Grant [grant no. 2018-67015-27495] and Hatch Project [accession no. 1019985] from the United States Department of Agriculture National Institute of Food and Agriculture.

## SUPPLEMENTARY TABLE LEGENDS

**Supplementary Table 1 |** Information on organisms in our dataset.

**Supplementary Table 2 |** Detailed information on organisms from the primary literature.

**Supplementary Table 3 |** Extended results for Fig. 4, showing all gene functions.

**Supplementary Table 4 |** Extended results for Fig. 4, showing all pathways.

**Supplementary Table 5 |** Examining gene functions of anaerobic organisms shows differences between fermentative and non-fermentative types.

**Supplementary Table 6 |** Examining MetaCyc pathways of anaerobic organisms shows differences between fermentative and non-fermentative types.

**Supplementary Table 7 |** Our reference metabolic model for predicting fermentation end products.

## SUPPLEMENTARY FIGURE LEGENDS

**Supplementary Fig. 1 | The definition of statistics we use to compare traits of fermentative and non-fermentative organisms. a**, Risk difference. **b**, Probability difference.

**Supplementary Fig. 2 | The log-odds ratio shows fermentation is more common in some phyla than others.** As Fig. 2b, except the effect size is measured using the log-odds ratio. Ca. = *Candidatus.*

**Supplementary Fig. 3 | Extended results for Fig. 3, showing category 2 isolation source.**

**Supplementary Fig. 4 | Extended results for Fig. 3, showing category 3 isolation source.**

**Supplementary Fig. 5 | Examining the raw data for cell length confirms that it differs between fermentative and non-fermentative organisms.**

**Supplementary Fig. 6 | Examining phenotypic traits of anaerobic organisms shows differences between fermentative and non-fermentative types. a**. Discrete traits. **b**, Continuous traits. Data are as Fig. 3, except only anaerobic organisms are included.

**Supplementary Fig. 7 | End products of fermentation previously reported for organisms of the human gut.** Data are from https://www.vmh.life/#microbes/fermcarb.

**Supplementary Fig. 8 | Reactions missing from organisms observed but not predicted to produce propionate or butyrate.** Adding any one of the missing reactions to the metabolic model restored the organism’s ability to produce this end product.

**Supplementary Fig. 9 | Good agreement exists between information in *Bergey’s Manual of Systematic of Archaea and Bacteria* and the primary literature. a**, Written descriptions (*n* = 88 organisms). Organisms are all type strains we found from one environment (the rumen). The year is from citations in Supplementary Table 1 and may differ from when an organism was originally isolated. **b**, Fermentative ability (*n* = 69 organisms). Organisms not in *Bergey’s Manual* are excluded. **c**, End products of fermentation (*n* = 52 organisms). Organisms with no fermentation products reported in *Bergey’s Manual* are excluded.

