## Supplementary Figures for "The phenotype and genotype of fermentative microbes"

Timothy J. Hackman✉
Department of Animal Science, University of California, Davis, CA, USA

✉

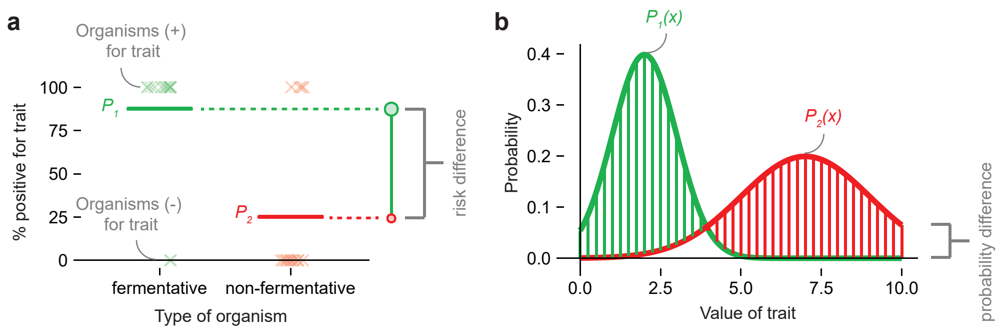


**Supplementary Fig. 1 | The definition of statistics we use to compare traits of fermentative and non-fermentative organisms.** **a**, Risk difference. **b**, Probability difference.

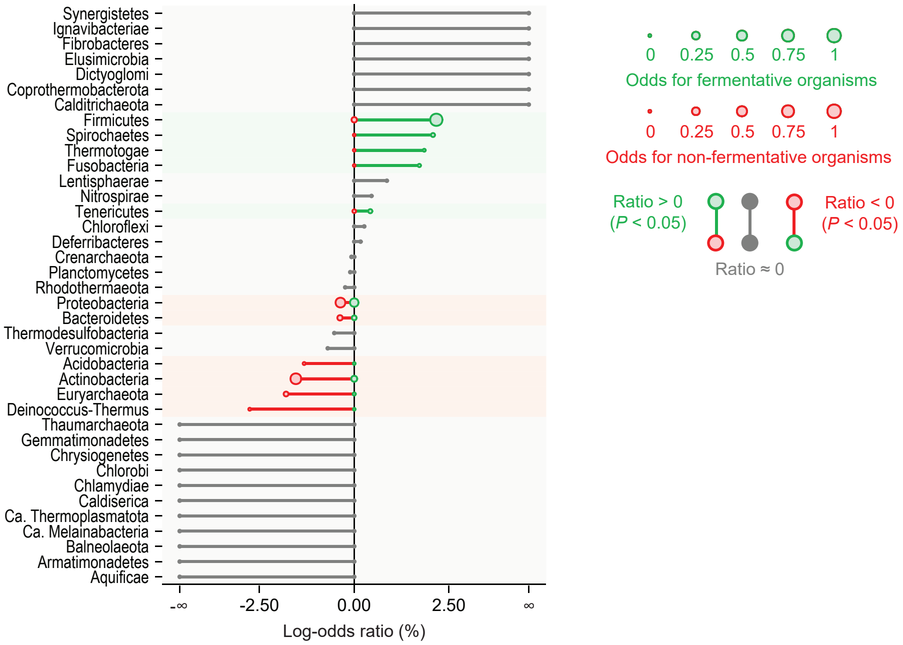


**Supplementary Fig. 2 | The log-odds ratio shows fermentation is more common in some phyla than others.** As Fig. 2b, except the effect size is measured using the log-odds ratio. Ca. = *Candidatus*.

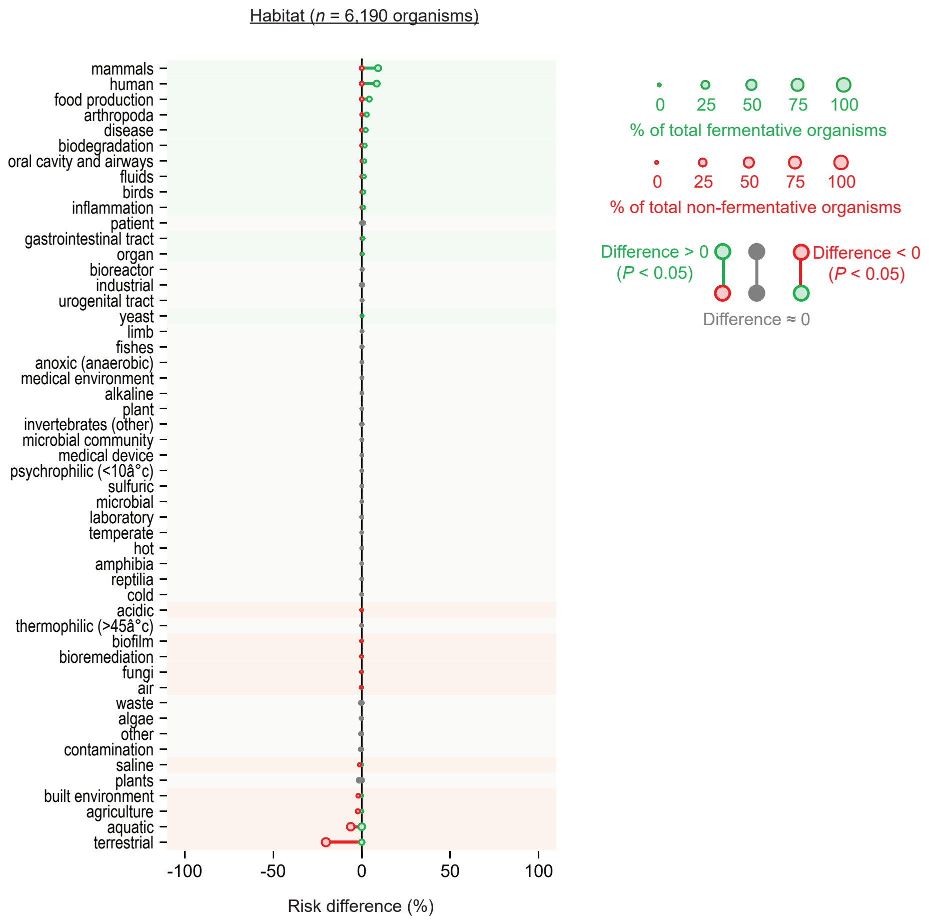


**Supplementary Fig. 3 | Extended results for Fig. 3, showing category 2 isolation source.**

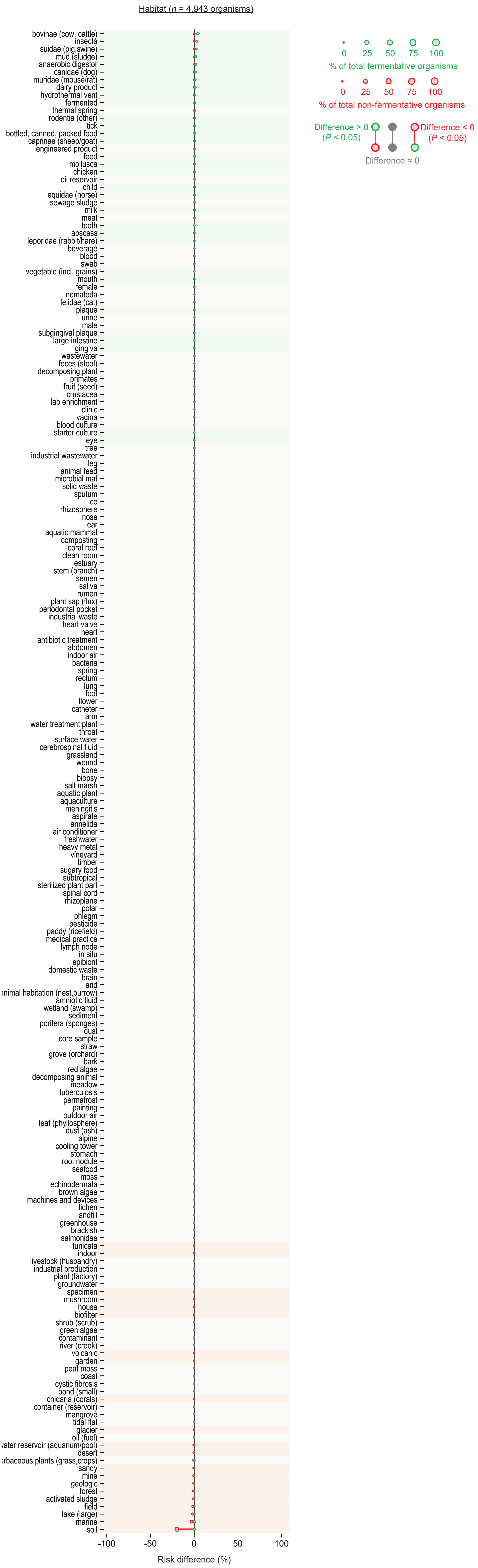

Supplementary Fig. 4 | Extended results for Fig. 3, showing category 3 isolation source.

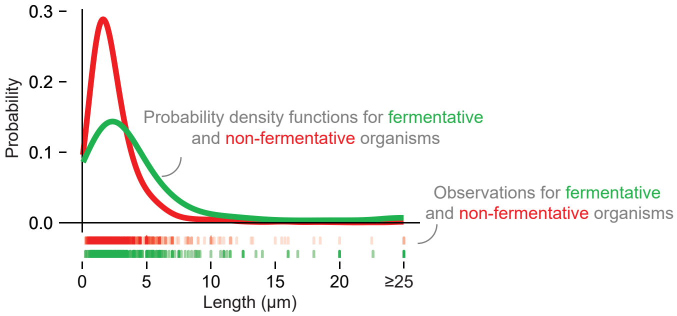


**Supplementary Fig. 5 | Examining the raw data for cell length confirms that it differs between fermentative and non-fermentative organisms.**

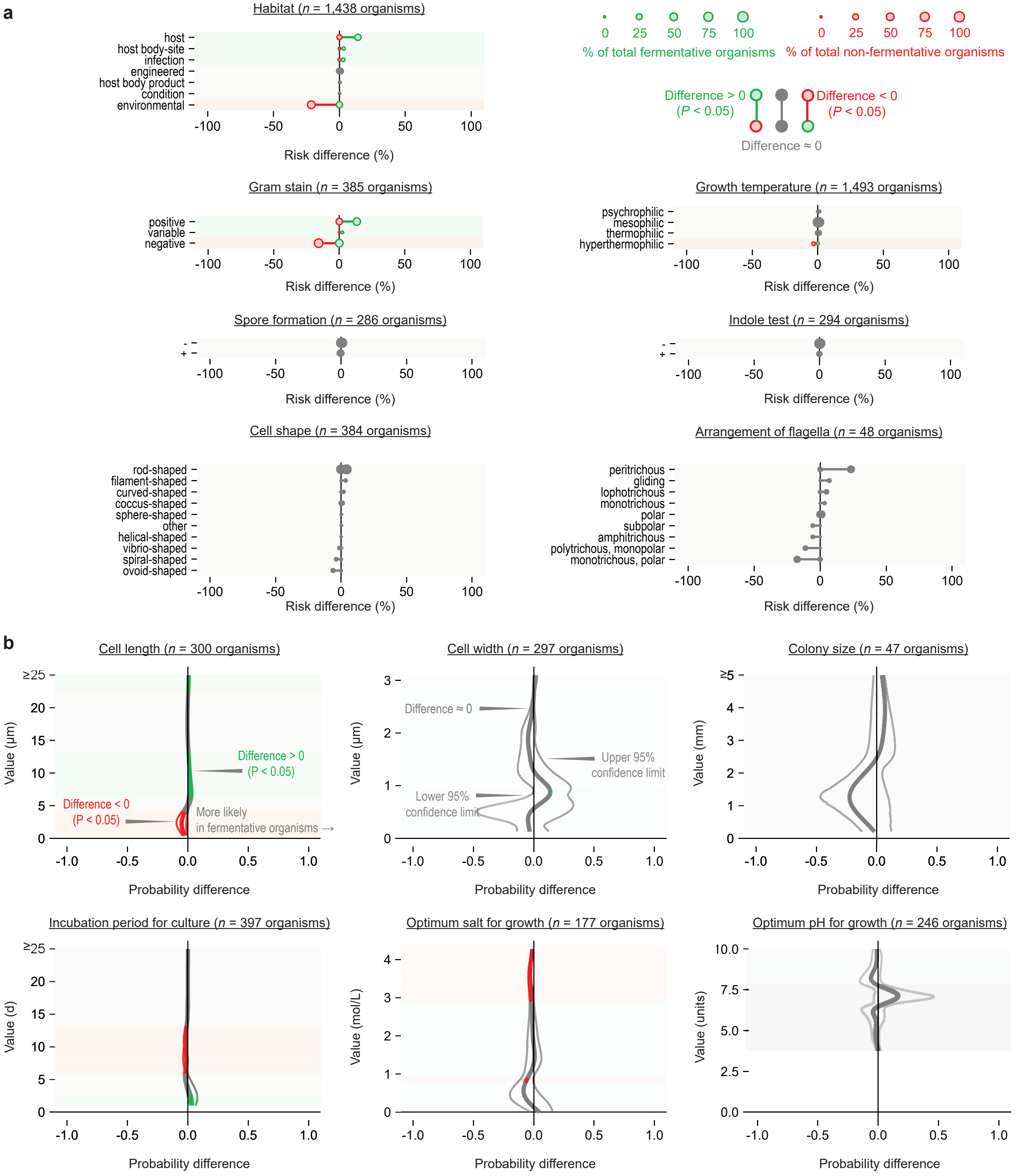


**Supplementary Fig. 6 | Examining phenotypic traits of anaerobic organisms shows differences between fermentative and non-fermentative types. a,** Discrete traits. **b,** Continuous traits. Data are as in Fig. 3, except only anaerobic organisms are included.

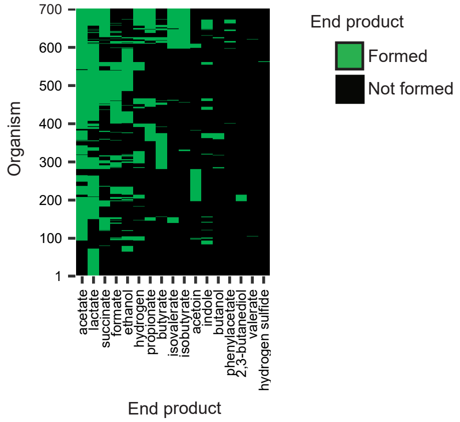


**Supplementary Fig. 7 | End products of fermentation previously reported for organisms of the human gut.** Data are from https://www.vmh.life/#microbes/fermcarb.

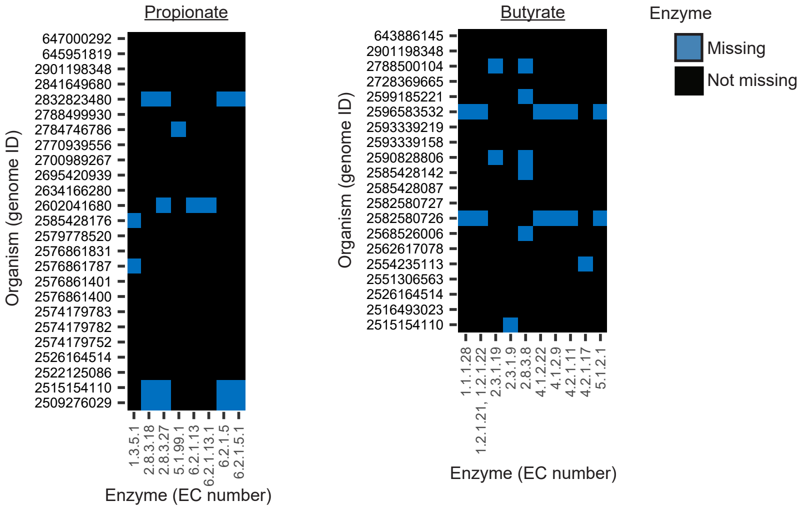


**Supplementary Fig. 8 | Reactions missing from organisms observed but not predicted to produce propionate or butyrate.** Adding any one of the missing reactions to the metabolic model restored the organism's ability to produce this end product.

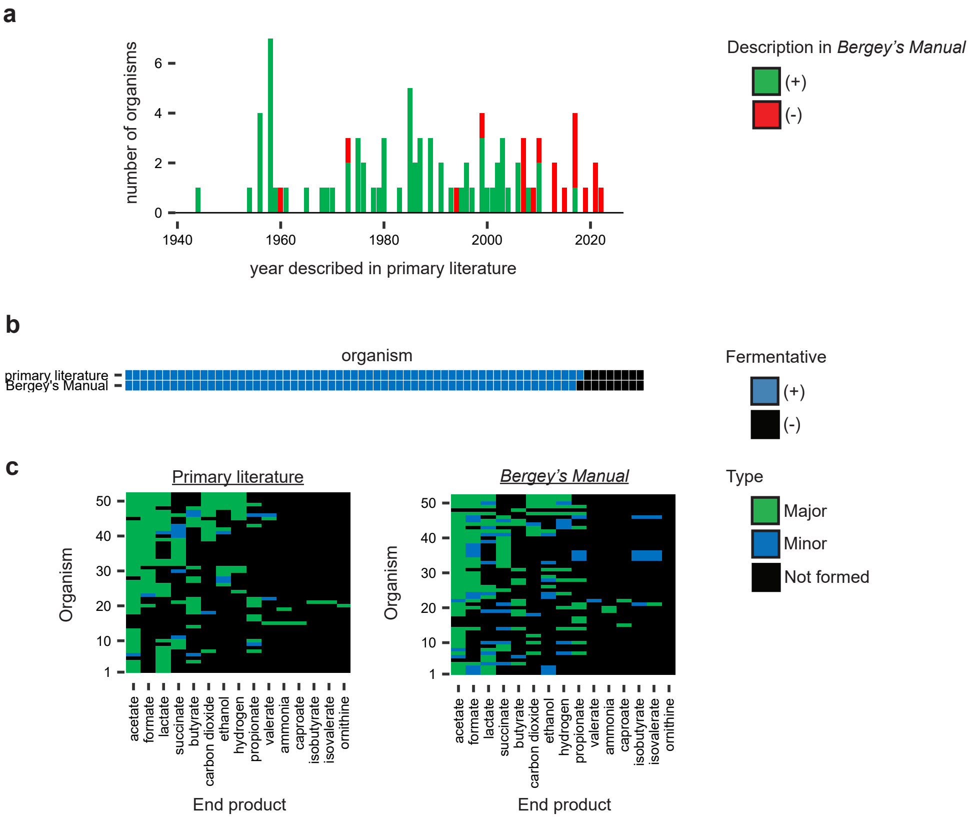


**Supplementary Fig. 9 | Good agreement exists between information in *Bergey's Manual of Systematic of Archaea and Bacteria* and the primary literature.** **a**, Written descriptions ($n$ = 88 organisms). Organisms are all type strains we found from one environment (the rumen). The year is from citations in Supplementary Table 1 and may differ from when an organism was originally isolated. **b**, Fermentative ability ($n$ = 69 organisms). Organisms not in *Bergey's Manual* are excluded. **c**, End products of fermentation ($n$ = 52 organisms). Organisms with no fermentation products reported in *Bergey's Manual* are excluded.
